## Supplemental figures and table for "Enhanced mitochondrial biogenesis promotes neuroprotection in human stem cell derived retinal ganglion cells of the central nervous system"

### Supplementary Figure 1

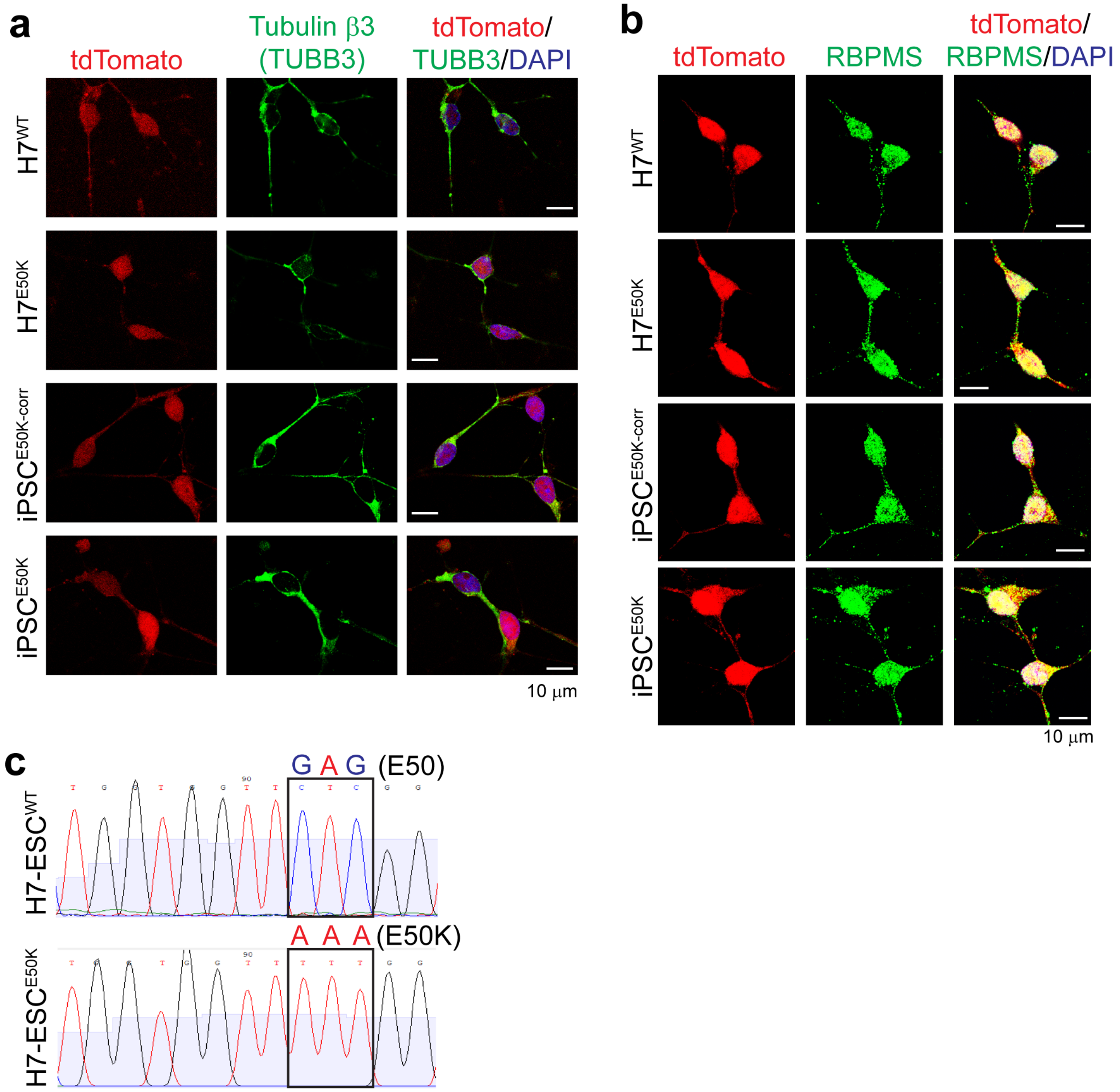


**Supplementary Figure 1. hRGC differentiation and CRISPR mutated OPTN$^{E50K}$ stem cell lines. (a-b)** Shown are confocal immunofluorescence images of hRGCs against **(a)** neuronal marker TUBB3 and **(b)** RGC specific marker RBPMS. Scale bars are 10 µm. **(c)** Sanger sequencing chromatograms show the E50 sequence and the homozygous E50K mutation in the H7-ESCs.

Supplementary Figure 2

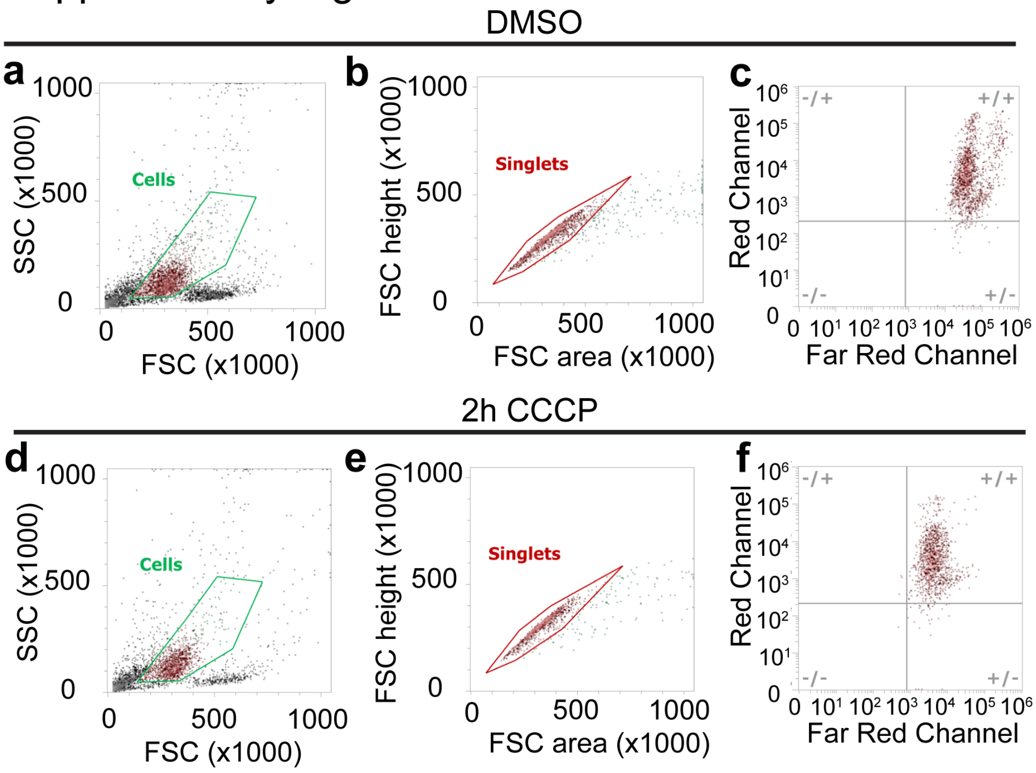


**Supplementary Figure 2. Flow cytometry measurement of MTDR labelled mitochondrial mass in hRGCs.** Live hRGCs were gated from the total population measured after **(a)** DMSO and **(d)** 2h 10 μM CCCP treatments. From the live cells, **(b, e)** singlet population was then gated, (**c, f)** subsequently analyzed for average MTDR intensity positive for both tdTomato (red) and MTDR (far red). **(f)** CCCP treatment reduces MTDR positive mitochondrial mass as shown by the left shift of MTDR positive cells.

### Supplementary Figure 3

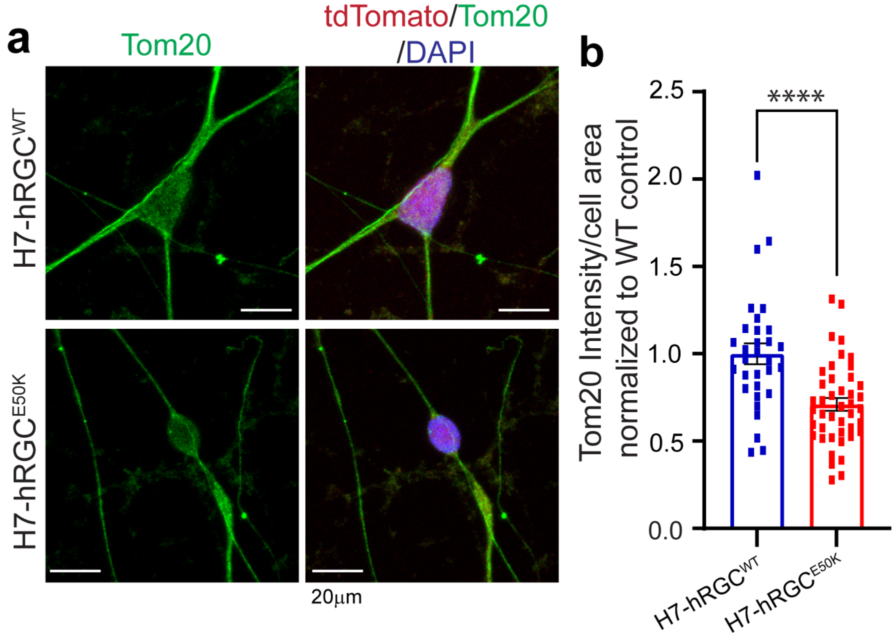


**Supplementary Figure 3. Glaucomatous *OPTN*$^{E50K}$ hRGCs possess less mitochondrial mass.** **(a)** Representative confocal immunofluorescence images of Tom20, tdTomato, and DAPI of untreated WT and E50K H7-hRGCs. **(b)** Quantification of Tom20 intensity from the sum projections of z-stacks relative to cell area, normalized to WT. Unpaired student's *t-test*, ****, p-value < 0.0001, n=30 (3 biological replicates).

### Supplementary Figure 4

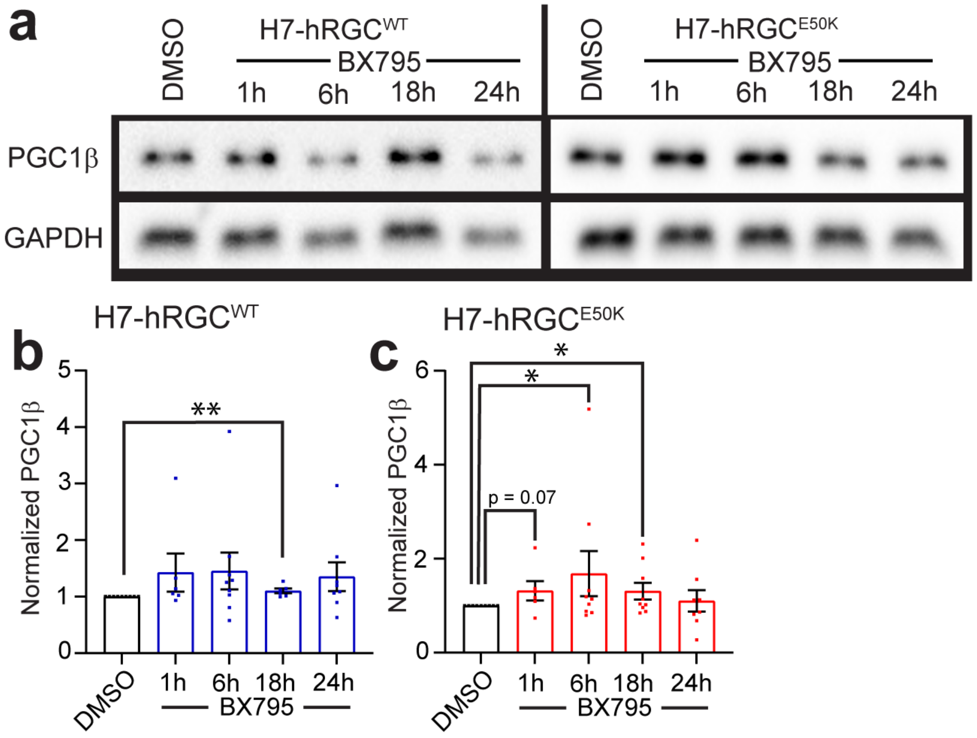


**Supplementary Figure 4. TBK1 inhibition by BX795 increases PGC1β level in hRGCs.** **(a)** Representative western blot images of H7-hRGCs treated with 1 μg/ml BX795 for the indicated timepoints. **(b, c)** Quantification of average western blot band intensities of PGC1β relative to its GAPDH loading control, then normalized to DMSO control, for **(b)** WT and **(c)** E50K hRGCs. n=3-9. Unpaired student's *t-test* between DMSO and the individual timepoints. * p-value < 0.05, ** p-value < 0.01.

#### Supplementary Figure 5

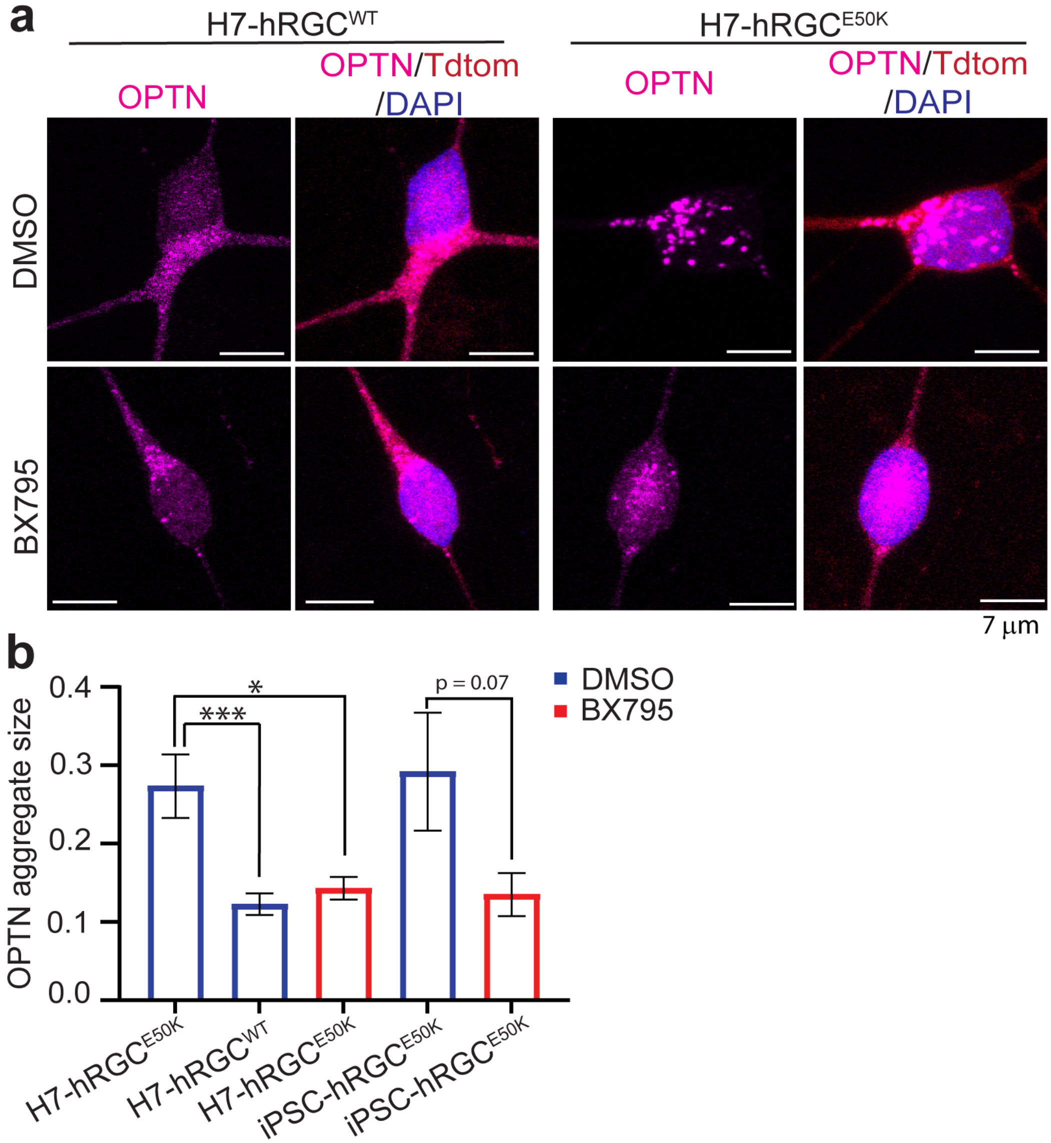


**Supplementary Figure 5. BX795 treatment dissolves *OPTN^E50K* aggregates in hRGCs.**
**(a)** Representative confocal immunofluorescence images of OPTN, tdTomato, and DAPI of H7-hRGCs treated with 1 µg/ml BX795 for 24hrs. **(b)** Quantification of OPTN aggregate size from the sum projections of confocal z-stacks. Mann Whitney U test between independent data sets * p-value <0.05, *** p-value <0.001, n=130-250 aggregates.

Supplementary Figure 6

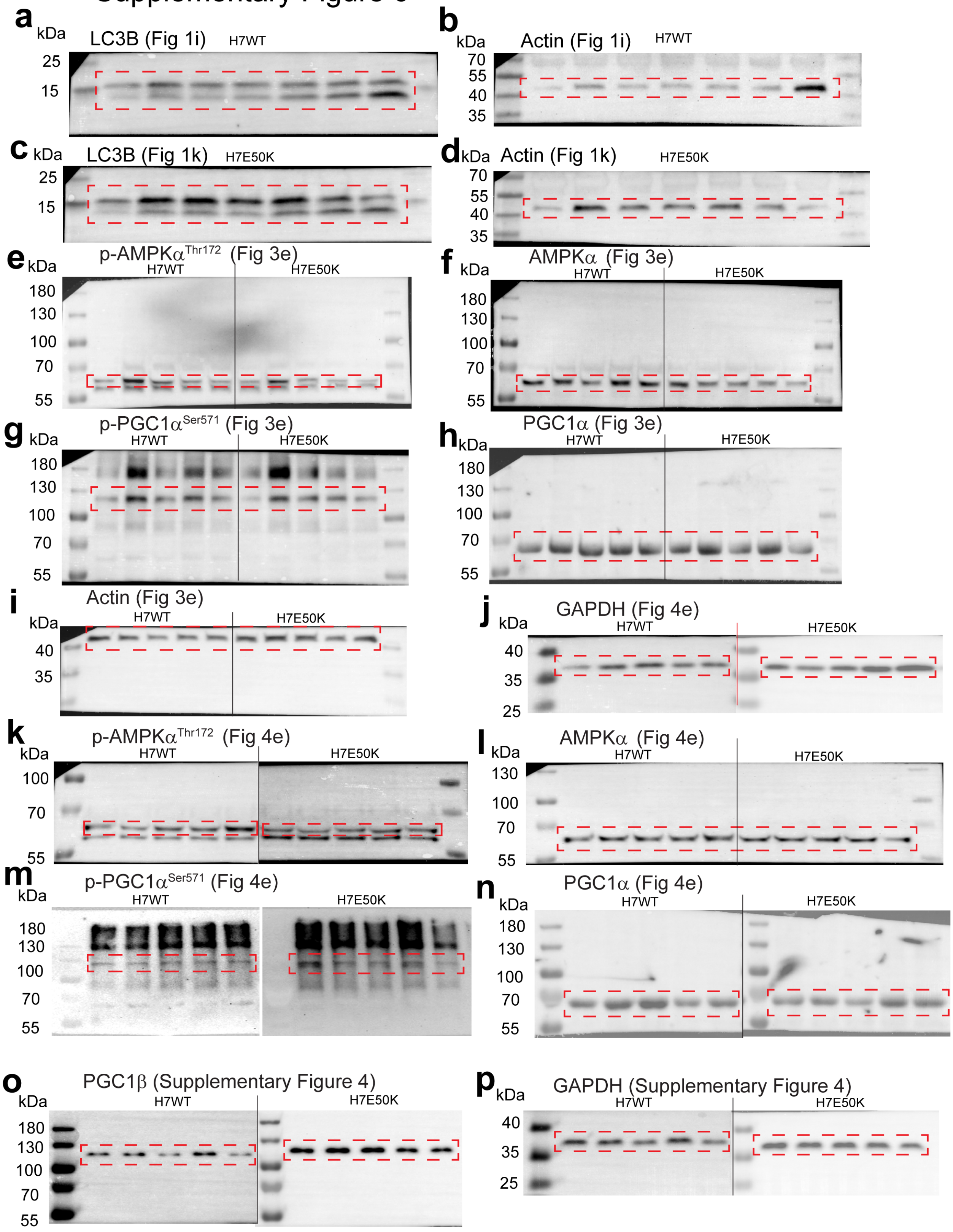

**Supplementary Figure 6. Full size images of representative western blots.**
Full blot images of western blot membranes shown in previous figures with molecular weight marker. Red boxes represent quantified protein bands.

**Supplemental table S1: list of reagents, software, antibodies, and primers (qPCR)**

| REAGENTS AND ASSAY KITS | | |
|---|---|---|
| **Reagent** | **Manufacturer** | **Catalog Number** |
| Matrigel | Corning | CB40230 |
| Gentle Cell Dissociation Reagent | Stem Cell Technology | 7174 |
| mTeSR1 | Stem Cell Technology | 85850 |
| Accutase | Sigma | A6964 |
| Blebbistatin | Sigma | B0560 |
| DMEM/F-12 | Thermo Scientific | 11330032 |
| Neurobasal media | Thermo Scientific | 21103049 |
| GlutaMAX supplement (100X) | Invitrogen | 35050061 |
| Antibiotic-Antimycotic | Thermo Scientific | 15240062 |
| B27 supplement | Thermo Scientific | 17504044 |
| N2 supplement | Thermo Scientific | 17502048 |
| Nicotinamide | Sigma | N3376 |
| IDE2 | Tocris | 4016 |
| Forskolin | Stem cell technologies | 3828S |
| Dorsomorphin | Stem cell technologies | P5499 |
| DAPT | Sigma | D5942 |
| MACS kit | Miltenyi Biotec | 130-091-051<br>130-042-108 |
| CD90.2 MicroBeads, mouse | Miltenyi Biotec | 130-121-278 |
| Dimethylsulphoxide (DMSO) | Sigma | 276855 |
| Carbonyl cyanide-3-chlorophenylhydrazone (CCCP) | Sigma | C2759 |
| BX-795 hydrochloride | Sigma | SML0694 |
| M-PER mammalian protein extraction reagent | Thermo Scientific | 78503 |
| Halt protease and phosphatase inhibitor cocktail (100X) | Thermo Scientific | 1861281 |
| 0.5 M EDTA solution (100X) | Thermo Scientific | 1861274 |
| DC Protein Assay Kit II | Bio-Rad | 5000112 |
| Laemmli buffer (4X) | Bio-Rad | 1610747 |
| PageRuler™ Prestained Protein Ladder | Thermo Scientific | PI26616 |
| BioRad Mini-PROTEAN TGX precast gels | Bio-Rad | 4561024<br>4561025 |
| Running buffer (10x Tris/Glycine/SDS) | Bio-Rad | 1610732 |
| Transfer buffer (10x Tris/Glycine) | Bio-Rad | 1610734 |
| Immuno-blot PVDF membrane | Bio-Rad | 1620260 |
| Methanol | Sigma | 34860 |
| TBS buffer 20x | VWR | J640 |
| Tween 20 | Sigma | P9416 |
| Clarity Max Western ECL substrate | Bio-Rad | 1705060 |
| DAPI | Molecular Probes | D1206 |
| MitoTracker™ Deep Red FM (MTDR) | Invitrogen | M22426 |
| Paraformaldehyde 16% solution, EM grade | Electron Microscopy Sciences | 15710 |
| Triton-X-100 | Sigma | T8787 |
| Donkey Serum | Sigma | D9663 |
| RNeasy Mini Kit | Qiagen | 74104 |

| | | |
|---|---|---|
| 5x all-in-one RT MasterMix (with AccuRTGenomic DNA Removal kit) | Applied biological materials | G492 |
| BlasTaq 2X qPCR MasterMix | Abcam | G891 |
| DNeasy Blood & Tissue Kit | Qiagen | 69506 |
| TaqMan Fast Universal PCR Master Mix (2X), no AmpErase UNG | Applied Biosystems | 4352042 |
| TaqMan™ Copy Number Reference Assay, human, RNase P | Applied Biosystems | 4403326 |
| JC1 dye | Cayman Chemicals | 10009172 |
| ApoTox-Glo™ Triplex Assay | Promega | G6320 |
| Seahorse XF Cell Mito Stress Test Kit | Agilent | 103015-100 |
| Seahorse XF DMEM medium, pH 7.4 | Aglient | 103575-100 |
| Seahorse XF 1.0 M glucose solution | Agilent | 103577-100 |
| Seahorse XF 100 mM pyruvate solution | Agilent | 103578-100 |
| Seahorse XF 200 mM glutamine solution | Agilent | 103579-100 |
| Oligomycin from *Streptomyces diastatochromogenes* | Sigma | O4876 |
| Antimycin A from *Streptomyces* sp. | Sigma | A8674 |
| Rotenone | Sigma | R8875 |
| 2-Deoxy-D-glucose | Sigma | D8375 |
| Carbonyl cyanide 4-(trifluoromethoxy)phenylhydrazone (FCCP) | Sigma | C2920 |
| **ANTIBODIES** | | |
| **Antibody** | **Manufacturer** | **Catalog Number** |
| β-actin (rabbit) | Cell signaling technologies | 4967S |
| GAPDH (rabbit monoclonal) | Cell signaling technologies | 2118S |
| PGC1α - N-terminal (rabbit) | Abcam | ab191838 |
| Human Phospho-PGC1α (S571) (rabbit) | R&D Systems | AF6650 |
| AMPKα (rabbit) | Cell signaling technologies | 2532S |
| Phospho-AMPKα (Thr172) (rabbit monoclonal) | Cell signaling technologies | 2535T |
| PGC1β (rabbit monoclonal) | Abcam | ab176328 |
| LC3B (rabbit) | Sigma | L7543 |
| Goat anti-rabbit-IgG1-HRP-linked Ab | Cell signaling technologies | 7074S |
| Optineurin (C-Term) (rabbit polyclonal) | Cayman Chemicals | 100000 |
| TOM 20 (mouse) | Santa Cruz | sc-17764 |
| Tubulin β 3 (TUBB3) (mouse) | Biolegend | 801202 |
| RBPMS (rabbit) | GeneTex | GTX118619 |
| Donkey anti-mouse Alexa Fluor 488 | Invitrogen | A12379 |
| Donkey anti-rabbit Alexa Fluor 647 | Invitrogen | A31573 |
| **SOFTWARE AND ALGORITHMS** | | |
| ImageJ | NIH | |
| Prism version 9 | GraphPad | |
| Zen Microscope Software | Zeiss | |
| Seahorse Wave Desktop | Agilent | |
| PrimerBank | https://pga.mgh.harvard.edu/primerbank/ | |

| **PRIMERS FOR qPCR** | | |
|---|---|---|
| GAPDH | PrimerBank ID: 83641890b1 | F: 5’AAGGTGAAGGTCGGAGTCAAC3’<br>R: 5’GGGGTCATTGATGGCAACAATA3’ |
| PGC 1α | PrimerBank ID: 116284374c1 | F: 5’TCTGAGTCTGTATGGAGTGACAT3’<br>R: 5’CCAAGTCGTTCACATCTAGTTCA3’ |

| | | |
|---|---|---|
| PGC 1β | PrimerBank ID: 289577089c1 | F: 5’GATGCCAGCGACTTTGACTC3’<br>R: 5’ACCCACGTCATCTTCAGGGA3’ |
| PRC | PrimerBank ID: 40807451c1 | F: 5’CAAGCGCCGTATGGGACTTT3’<br>R: 5’GGAGGCATCCATGTAGCTCT3’ |
| NRF 1 | PrimerBank ID: 93141038c1 | F: 5’AGGAACACGGAGTGACCCAA3’<br>R: 5’TATGCTCGGTGTAAGTAGCCA3’ |
| NRF 2 | PrimerBank ID: 372620347c1 | F: 5’TCAGCGACGGAAAGAGTATGA3’<br>R: 5’CCACTGGTTTCTGACTGGATGT3’ |
| ND1 F | https://doi.org/10.1007/s13277-014-2937-2 | 5’CCTTCGCTGACGCCATAAA3’ |
| ND1 R | | 5’TGGTAGATGTGGCGGGTTTT3’ |
| ND1 Probe | | 6FAM-5’TCTTCACCAAAGAGCC3’-MGBNFQ |
